# From Images and Words to Modality-Independent Semantic Representations: When and Where They Emerge

**DOI:** 10.64898/2026.10.30.763035

**Authors:** Nicolas Piron, Théo Desbordes, Itsaso Olasagasti, Sophie Schwartz, Nina Kazanina

**Affiliations:** Department of Basic Neurosciences, Faculty of Medicine, University of Geneva, Switzerland; Swiss Center for Affective Sciences, University of Geneva, Geneva, Switzerland

## Abstract

Conceptual knowledge can be accessed from different modalities, such as seeing an image or reading a written word. While prior neuroimaging studies implicate anterior temporal cortex in modality-independent semantic representations, when and how representations evoked by images and words converge onto a modality-independent code remains unclear. Moreover, evidence for modality-independent category representations comes largely from tasks that explicitly required semantic processing, leaving open whether such representations emerge when semantic category and stimulus modality are task-irrelevant. We magnetoencephalography (MEG) to determine when and where modality-independent semantic category representations emerge across images and words when participants (n = 32) encoded six-item sequences containing an exemplar of six semantic categories counterbalanced across positions and reported the ordinal position of a probed item. Decoding performance was above chance for all categories in each modality from approximately 70 ms relative to item presentation. Generalization peaked at similar latencies for the two modalities (∼520-530 ms), where image and word classifiers had the most similar coefficients. In both transfer directions, activity elicited by images generalized mostly to later activity elicited by words, and a given period of the image response shared its category code with a longer period of the word response. Source-space decoding around the peak of generalization identified the same regions for both transfer directions: left and right anterior temporal lobe, perisylvian cortex (more extensive in the left hemisphere), right visual cortex, and bilateral orbitofrontal cortex, indicating a modality-independent category code distributed across the cortex.

## Introduction

A concept can be activated by a photograph or a printed word, accessing the same underlying knowledge. Thus, concepts may involve neural representations that are largely independent of the format of the input. How the brain represents conceptual knowledge through neural representations that are independent of input modality remains an open question [1,2]. Neuropsychological and functional MRI (fMRI) research suggests that the anterior temporal lobe may function as a hub where information arriving through modality-specific networks is recoded into an amodal conceptual representation [3,4] that is recruited during semantic processing irrespective of input modality [5,6]. Format-independent semantic information has also been demonstrated in temporal, parietal and prefrontal cortex [7,8]. What remains unclear is when the representations evoked by images and by written words converge onto a modality-independent code, and how this convergence unfolds across brain regions. Addressing both questions requires a time-resolved measure of representational content that enables direct comparison across input modalities and localization in the brain.

Cross-modal classification of MEG data suits well this purpose. In decoding analysis, a classifier is trained on the activity distributed across sensors; training a classifier in one context and testing it in another has become a standard way of establishing that the *discriminative structure* is shared between the two contexts [9–11]. Indeed, two modalities may elicit covarying neural signal, for instance if both drive a given region more strongly for animate than for inanimate items, without a classifier trained on one modality being able to read out the other. Successful cross-modal decoding is a stronger claim: it requires the geometry of the representation, that is, the direction along which the categories are separated, to be common to both modalities. Applied to semantic categories presented as images and as written words, above-chance cross-modal classification thus indicates that a representation common to the two formats is expressed. Coupled with time-resolved neuroimaging such as MEG, cross-modal decoders can be applied millisecond by millisecond, rather than over the timescale of seconds captured by the haemodynamic response in fMRI. Fitting and testing classifiers at every pair of samples evaluates each combination of a training and a testing time independently [12]. Training on one modality (e.g. images) and testing on the other (e.g. words) then recovers whether a shared format of representation exists, when it first emerges, and how long it persists.

Several MEG studies tested whether semantic category information generalizes between images and written words [13–16]. With the exception of the naming-aloud study by Dirani and Pylkkänen [14], if cross-modal generalization was observed, the modality-independent representation emerged earlier for images than for words, consistent with more direct access to category information from an image than from a written word. In all these cases, however, the task made category membership or the correspondence between an image and a word explicit, through naming and reading [14], matching the items between modalities [13], or semantic judgment [15,16]. It is therefore unknown whether a modality-independent representation, and the earlier availability of category information from images, emerge spontaneously when the task makes no reference to category or modality.

Although some of these MEG studies investigated the spatial distribution of cross-modal generalization at the sensor level [13,14,16], to our knowledge none has localized time-resolved cross-modal generalization in source space. Thus, the cortical network supporting cross-modal generalization remains undetermined.

We used an MEG classification and generalization approach followed by source-localization to determine when and where modality-independent semantic category representations emerge across images and words in a task that makes no reference to semantic information. Participants viewed sequences of six items, either images or words, drawn from six semantic categories; their task was to identify the ordinal position of a probed item in the sequence and thus did not direct attention to either its semantic category or its modality. Crucially, category was orthogonal to position and therefore irrelevant to the task. We decoded the six semantic categories with one-versus-rest classifiers within each modality, applied them to the other modality, and compared the geometry of the decision hyperplanes across modalities. Finally, to localize the cortical sources of cross-modal category information, we projected single-trial data into source space and repeated the decoding on the cortical surface with a searchlight, that is, separately within a small patch of cortex centred on each vertex. Classifiers were trained on one modality and tested on the other, using a time window centred on the peak of temporal cross-modal generalization.

We hypothesized that (i) cross-modal generalization emerges in a task without any explicit reference to category membership or cross-modal matching, (ii) modality-independent semantic (here category) representation is reached earlier from images than from written words, and (iii) the anterior temporal lobe is implicated in generating modality-independent representations.

## Methods

### Participants

Thirty-two participants took part in the study (21 female, mean age = 23.5 years, SD = ± 2.7). The sample size was chosen to match previous studies that employed decoding for cross-modal analyses with MEG [13,14]. All participants self-reported being right-handed, having normal or corrected-to-normal vision, and no history of neurological disorders. The study was conducted at Campus Biotech, Geneva, and was approved by the Geneva’s cantonal ethics committee. All participants gave written informed consent and were compensated at the rate of 30 CHF/hr, with a small bonus for performance. No participants were excluded from the analyses.

### Data acquisition

MEG data were recorded using a MEGIN Triux Neo system equipped with 306 SQUID sensors comprising 204 planar gradiometers and 102 magnetometers and a sampling rate of 1000 Hz. Participants were seated upright. Prior to each recording session, head shape was digitized using a Polhemus FASTRAK system, including three anatomical landmarks (nasion, left and right preauricular points) and additional head shape points recorded in string mode. Five continuous head position indicator (cHPI) coils were used to track head position continuously during recording. Auxiliary channels included electrooculography (horizontal and vertical), electrocardiography, and a photodiode channel registering stimulus onset timing.

Visual stimuli were backprojected onto a 55.0 × 30.5 cm translucent screen positioned at 120 cm from the participant’s eyes using a ProPixx projector set to 25% luminosity and auditory stimuli were delivered binaurally via MEG-compatible pneumatic earphones and SoundPixx system (VPixx Technologies, Canada). The volume was adjusted for each participant based on their perceptual threshold (chosen values ranging from-55 to-24 dBFS). A photodiode attached to the screen recorded the precise onset of the visual stimuli with respect to the MEG recording. Participants responded using a 4-button response pad. Following the MEG session, a structural T1-weighted MRI scan was acquired on a 3T Siemens MAGNETOM Prisma scanner, to construct an individual head model for MEG source localization.

### Stimuli and task design

The stimuli consisted of 72 images and 72 corresponding French words, drawn from six semantic categories: animals, body parts, cartoon characters, colours, landscapes and shapes (12 exemplars per category). Each category featured a consistent visual style (either in colour or black-and-white) for images or font (always black) for words (see Supplementary Material for details). The rationale was two-fold: a uniform visual style within each category was expected to reinforce category-level representations for the participants, as well as homogenize early sensory representations and thus reinforce classification based on low-level visual features. Word length ranged from 3 to 9 letters. Two days before the MEG session, participants completed an online familiarization task during which they were exposed to every image-word pairs once.

The stimulus material was organized into 12 sequences of 6 items each, each sequence containing exactly one exemplar per category. Each sequence was defined by its items and by its category order, and was associated with a specific note, played by an instrument specific to that sequence, which accompanied the onset of each of the 6 items during presentation (no notes were played during probe phases). Every sequence was presented in both modalities during the experiment (Fig. 1a).

**Figure 1.**
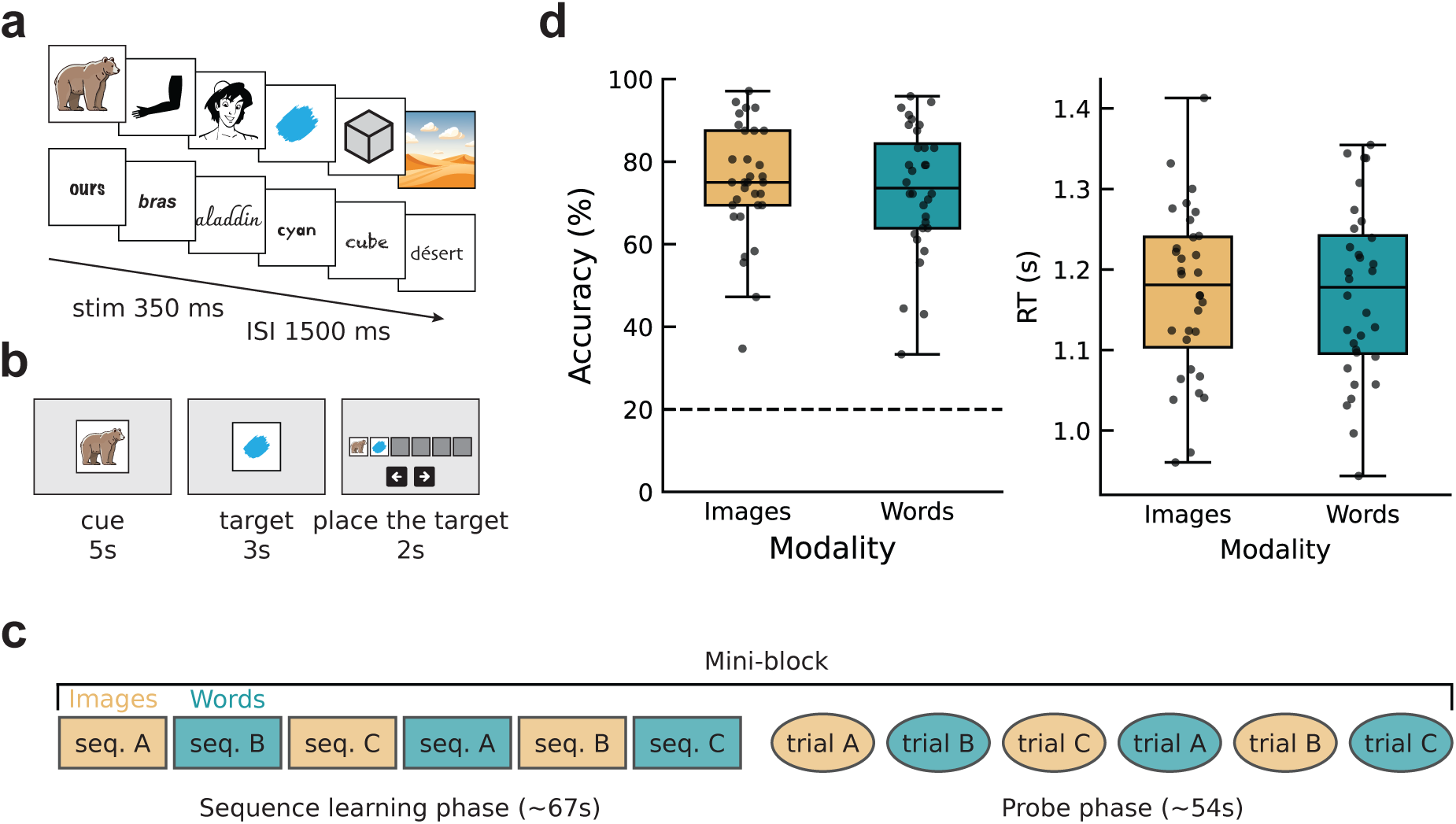
Experimental design and behavioural data. (a) Example sequence with images (top) and words (bottom). Stimuli were presented for 350 ms, with an interstimulus interval of 1500 ± 50 ms. (b) Example test trial. Participants first saw a cue, always the first element of a sequence, followed by the target. They then had to place the target at its correct position within the sequence with arrow keys. (c) Mini-block design. Three sequences (seq. A, seq. B, seq. C) are each presented twice, once as images and once as words, with modality alternating from one presentation to the next (left). Six test trials, one per sequence and modality (right). Colour indicates modality: yellow for images, blue for words. (d) Behavioural accuracy (left) and reaction time (right) for the sequence position question, split by stimulus modality (images or words). Each dot represents one participant, and the dashed line indicates chance level (1/5). Paired t-tests revealed no significant difference between modalities. All participants performed above chance level.

The experiment was organized into mini-blocks (Fig. 1c). Each mini-block presented three of the twelve sequences, once as images and once as words, with modality alternating from one presentation to the next: sequence A as images, B as words, C as images, then A as words, B as images, C as words. A mini-block therefore comprised 36 items (6 sequence presentations × 6 items), followed by a probe phase of six trials. Items appeared one at a time for 350 ms, separated by an inter-stimulus interval of 1500 ± 50 ms, with no additional delay between the last item of one sequence and the first item of the next (Fig. 1a).

Participants completed 24 mini-blocks over two recording runs, with a break after every three mini-blocks. In the second run, the category orders of the first run were reused with entirely different exemplars, constituting 6 new sequences (12 in total). Each sequence therefore appeared in six mini-blocks and was presented six times as images and six times as words, yielding 72 presentations per category and per modality. Stimuli were presented with PsychoPy v2024.1.4. [17].

At the end of each mini-block, participants completed a probe phase composed of six trials, one per sequence-modality combination presented during the mini-block (Fig. 1b). Each probe trial began with a 5 s-long, gradually fading out cue (image or word), always corresponding to the first item of a sequence and thus cuing the probed sequence. A target item, drawn randomly from the remaining five items, then appeared for a recall period of 3 s. Participants were asked to report the position of the target item in the sequence using a button interface displaying six slots. The cue was placed in the first slot and the target initially in the second; participants moved the target one slot to the right or to the left with a button press and confirmed its final position with a separate button.

### MEG data preprocessing

MEG data were preprocessed using MNE-Python v1.12 [18]. Bad channels were identified through visual inspection (on average, 2.4 channels per participant) and interpolated during the subsequent Maxwell filtering step. Head position was estimated continuously throughout each recording from the cHPI coils, and a time-weighted average head position was computed. Signal Space Separation (Maxwell filtering; Taulu & Simola, 2006) was then applied to each recording with movement compensation, using the time-weighted average position as the destination, together with site-specific cross-talk correction and fine calibration. The data were band-pass filtered between 0.1 and 200 Hz, notch filtered at 50, 100 and 150 Hz to remove power line noise and its harmonics and downsampled to 500 Hz. Independent Component Analysis (ICA) was fitted on the concatenated recordings within a session, excluding the empty-room recording; retaining enough components to account for 99% of the variance. Components corresponding to ocular and cardiac artefacts were identified automatically from the EOG and ECG channels respectively and further confirmed by visual inspection prior to exclusion. The resulting ICA solution was applied to the MEG data.

After realignment of event triggers to stimulus onsets using the photodiode signal, the continuous data were segmented into epochs time-locked to the onset of each image or word during the sequence presentation phase (−200 to 1000 ms); epochs from the probe phase were not included. Epochs with peak-to-peak amplitude exceeding 7000 fT in any magnetometer were rejected automatically, resulting in an average rejection rate of less than 1%. Epochs were subsequently downsampled to 100 Hz prior to decoding.

### Temporal decoding models

For each category, an L2-regularized logistic regression classifier (scikit-learn v1.8 implementation, with balanced class weights and “liblinear” solver; Pedregosa et al., 2011) was trained using a one-versus-rest (OVR) scheme whereby a separate binary classifier was trained to distinguish each category from the remaining five. This approach yielded a distinct decoding time course for each of the six semantic categories, extending cross-modal generalization beyond the binary contrasts examined in previous studies. A linear classifier was used because its weights can be examined to identify the sensors that contribute to the decoding. Unless stated otherwise, decoding performance is reported as the area under the ROC curve (AUC) averaged across the six category-specific classifiers, referred hereinafter as ‘class-average’. The regularization strength C was selected on a subset of five participants prior to the main analysis by comparing decoding performance across a logarithmically spaced range of candidate values (0.001 to 10), with C = 0.01 yielding the best performance. This value was then fixed for all time points and all participants. Decoding was performed independently at each time point using 50-ms overlapping windows (five samples: the current sample and the four preceding it), by concatenating the sensor data across each window into a single feature vector. All further processing was applied to the feature vectors of each window separately, with parameters estimated on the training trials only and then applied to the test trials. Each feature was centered on its median and scaled by its interquartile range (25th to 75th percentile) across training trials. Dimensionality was then reduced with principal component analysis (PCA), retaining the components explaining 97% of the variance. To avoid border effects associated with the windowed feature extraction, the first 100 ms of the resulting AUC time courses and activation patterns were discarded, restricting the reported results to −100 to 1000 ms relative to stimulus onset.

### Temporal within-modality decoding

Decoding performance was first assessed separately within each stimulus modality (images, words) using 5-fold stratified cross-validation. For each fold, the classifier described above was trained and tested at each time point, and performance was quantified as AUC, averaged across folds. To characterize the temporal dynamics of these representations, classifiers trained at each time point were additionally tested at every other time point, yielding a train-time by test-time generalization matrix of AUC values for each modality [12].

### Temporal cross-modal decoding

To assess whether category information generalized across stimulus modalities, classifiers were trained on all data from one modality and tested on all data from the other, at every combination of training and testing time points, yielding a train-time by test-time generalization matrix of AUC values. This was performed in both directions, training on images and testing on words, and training on words and testing on images. In both cases, the training time is represented on the y axis and the testing time on the x axis (Fig. 2a, b).

**Figure 2.**
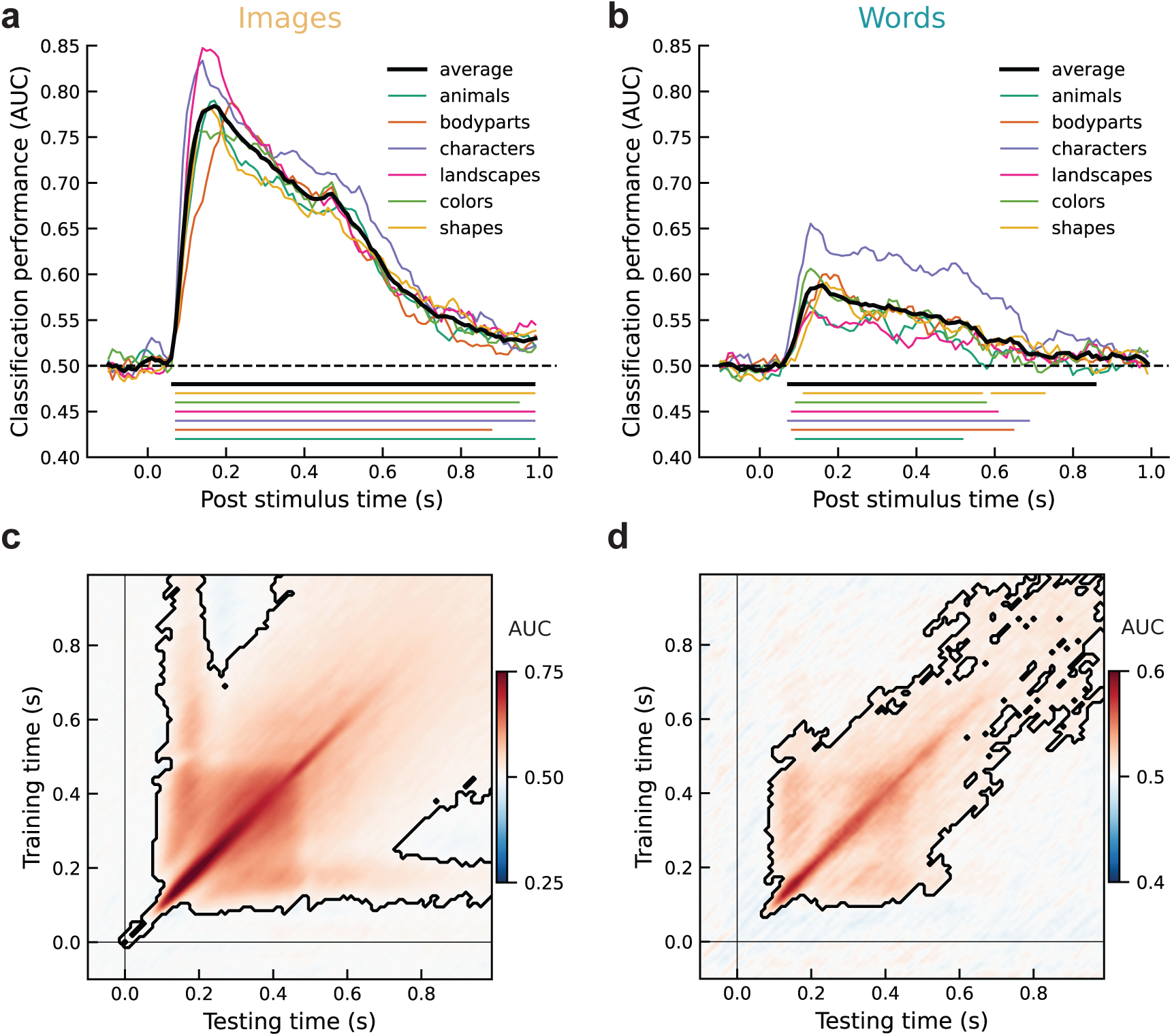
Decoding semantic category within modality. (a, b) One-versus-rest classification performance (AUC) as a function of time, for images (a) and words (b). Coloured lines show individual category decoding; the black line shows the average across categories. Horizontal lines below each panel indicate significant clusters (cluster-based permutation test) for the average and for each category separately. (c, d) Temporal generalization matrices for the class-averaged decoder, for images (c) and words (d). Colour indicates classification performance (AUC) for classifiers trained at one time point (y-axis) and tested at another (x-axis). Black outlines indicate significant clusters (cluster-based permutation test). Thin black lines mark stimulus onset.

### Decision subspace angle

Decoding performance describes how well two categories are separated, but not how the classifier achieves the separation: the same performance can be obtained by weighting the sensors in quite different ways. Because performance reduces a whole coefficient vector to a single scalar, it discards the geometry of the solution. We therefore complemented it with the Rotation Angle [21], which compares two classifiers by the direction of their coefficient vectors rather than by their accuracy. Because correlated noise can inflate weights on sensors carrying no signal, we used activation patterns rather than raw weights [22]. The resulting coefficient vector is referred to as the decision subspace.

For each pair of classifiers, the Rotation Angle θ was obtained as the arc cosine of the cosine similarity between their coefficient vectors. θ = 0° indicates perfectly aligned subspaces, θ = 90° orthogonal ones and θ = 180° perfectly reversed ones; for unrelated coefficient vectors, θ ≈ 90° is expected, which provides the reference against which observed angles are read. Because cosine similarity is normalized by the vector norms, θ depends only on the relative distribution of weight across sensors and not on the overall amplitude of either pattern and is therefore comparable between conditions that differ in decodability strength. Following Desbordes et al. [21], angles were computed on the subset of sensors contributing significantly to decoding (the union across time points of sensors whose absolute coefficients exceeded their pre-stimulus baseline (one-sample t-tests, p < 0.001, FDR-corrected), retaining 72 of 306 sensors) to avoid dilution of the angle measure from coefficients that predominantly reflect noise. Angles were computed separately for each category and averaged. The procedure was applied within each modality, yielding a symmetric matrix with a zero diagonal by construction, and across modalities, yielding a matrix whose axes correspond to image time (y) and word time (x) and which is neither symmetric nor zero on the diagonal, since it compares two distinct sets of classifiers.

### Cross-modal decoding in source space

To estimate which cortical regions supports category information, decoding was repeated in source space within a fixed time window, yielding one decoding score per cortical location (for similar approaches, see Krogh and Pylkkänen [23]; Zuanazzi et al. [24]. The window was a 100 ms × 100 ms train-test tile centred on the peak of the cross-modal generalization, defined as the time point pair with the highest decoding score (see Results, Cross-modal generalization).

Individual cortical surfaces were reconstructed from the structural MRI using the FreeSurfer recon-all pipeline, as implemented in MNE-Python. A boundary element model was built with the watershed algorithm, with three layers (inner skull, outer skull, scalp) and conductivities of 0.3, 0.006 and 0.3 S/m. The MEG-MRI coregistration was based on fiducial landmarks and digitized head points. Source space was an oct-5 subdivision of the white matter surface (1026 vertices per hemisphere). The noise covariance matrix was estimated from empty-room recordings preprocessed like the task data. The inverse operator used a loose orientation constraint (0.2) and depth weighting (0.8). Single-trial MEG data were projected onto the cortical surface with eLORETA (λ² = 1/9), and the component normal to the cortical surface was retained. Vertices of the medial wall (FreeSurfer “unknown” label) were excluded.

Decoding was performed with a surface searchlight, using as features the source time courses of each vertex and of its direct neighbours on the cortical mesh (6.9 vertices on average; mean radius 11.8 mm) within the training or testing window. The classifiers were the same as in the sensor-space analysis, without PCA. Classifiers were trained on one modality and tested on the other, in both directions, yielding one AUC per vertex and category, then averaged across categories for each participant.

## Statistical analysis

Statistical significance of decoding performance was assessed using cluster-based permutation testing [25,26], which corrects for multiple comparisons across time points without assuming independence between them. For each temporal analysis (time-resolved AUC and time generalization matrices), a one-sample, one-tailed t-test was computed at each time point, or pair of time points, testing the group-level average AUC against chance level (0.5). Adjacent time points, or pairs of time points, exceeding a cluster-forming threshold were grouped into clusters. This threshold was set to the t value corresponding to an uncorrected p of 0.05, one-tailed (t(31) = 1.70). The sum of t values within each cluster was compared against a null distribution obtained by repeating the procedure 10,000 times with randomly sign-flipped data, and clusters with a Monte Carlo p value below 0.01 were considered significant. The same procedure was applied to the source-space AUC maps (averaged across categories), with clusters formed over vertices instead of time points. The maps were morphed to the fsaverage surface, and spatial adjacency was defined from its source space. The cluster-forming threshold was p = 0.001. Clusters were retained if their Monte Carlo p value was below 0.01, a more stringent criterion than for the time-resolved analyses, adopted to restrict the reported maps to the most spatially extended effects. After statistical testing, we assigned the significant vertices to regions of the Destrieux atlas (aparc.a2009s [27])to interpret the source-space results anatomically.

For the cross-modal generalization matrices, we assessed the offset in the distribution of significant generalization pairs of time-points with respect to the diagonal. This allowed us to quantify whether one modality would tend to reach a neural state or representation earlier or later than the other modality. We defined the diagonal offset index as

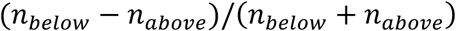

where *n_belo_*_w_ and *n_above_* are the numbers of significant time-point pairs falling below and above the diagonal respectively. A negative index indicates that representations at earlier training times generalize to later testing times, and a positive index the reverse. This index is a descriptive summary of the distribution of the significant cluster relative to the diagonal rather than a quantity subjected to statistical testing.

## Data availability

Behavioural and preprocessed MEG data will be available on OpenNeuro (openneuro.org/datasets/ds009022) upon publishing.

## Code availability

Code for the decoding and cross-modal analyses is available on GitHub (https://github.com/NicolasPiron/xmod-generalization).

## Results

### Behavioural data

Participants performed the sequence position task above chance level for both images (p < 0.001) and words (p < 0.001; Fig. 1d). Mean accuracy did not significantly differ between modalities (images: M = 74.9%, SD = ± 13.9; words: M = 73.3%, SD = ± 15.3; paired t-test, p = 0.125), nor did reaction times (images: M = 1.17 s, SD = ± 0.10; words: M = 1.17 s, SD = ± 0.11; p = 0.898).

### Within-modality category decoding

We first aimed to decode the six semantic categories from MEG activity separately for images and words, using logistic regression and one-versus-rest classification (Fig. 2a, b). For images, the class-average reached significance from 60 to 1000 ms, peaking at 0.78 AUC at 160 ms. For words, the decoding performance was significant from 70 to 860 ms, with the class-averaged decoder peaking at 0.59 AUC at 160 ms. A lower decoding performance for words than for images is in line with previous reports [11,28,29]. Early decoding in both modalities is expected to draw on low-level features, the shared visual style within each image category and the category-specific font for words, consistent with the stimulus design. The class-average temporal generalization profiles differed between modalities. For images, the significant cluster extended broadly along and around the diagonal from approximately 70 to 1000 ms, with classifiers trained around 100 ms partially generalizing until the end of the epoch, indicating that part of the early representation was maintained throughout. For words, the cluster likewise extended beyond the diagonal, but less uniformly: generalization was strongest early, with classifiers trained around 150 ms generalizing to approximately 600 ms and weakened at later training times, indicating that the representation evolved over the epoch.

### Cross-modal generalization

To assess whether the neural representation of a semantic category is shared across modalities, classifiers trained on one modality were tested on the other one. In both directions, significant clusters of above-chance generalization were obtained in the class-average approximately 240 ms to 1000 ms (Fig. 3a, b), indicating that the category representation included a modality-independent component. Generalization matrices for individual semantic categories are shown in Supplementary Figure 1.

**Figure 3.**
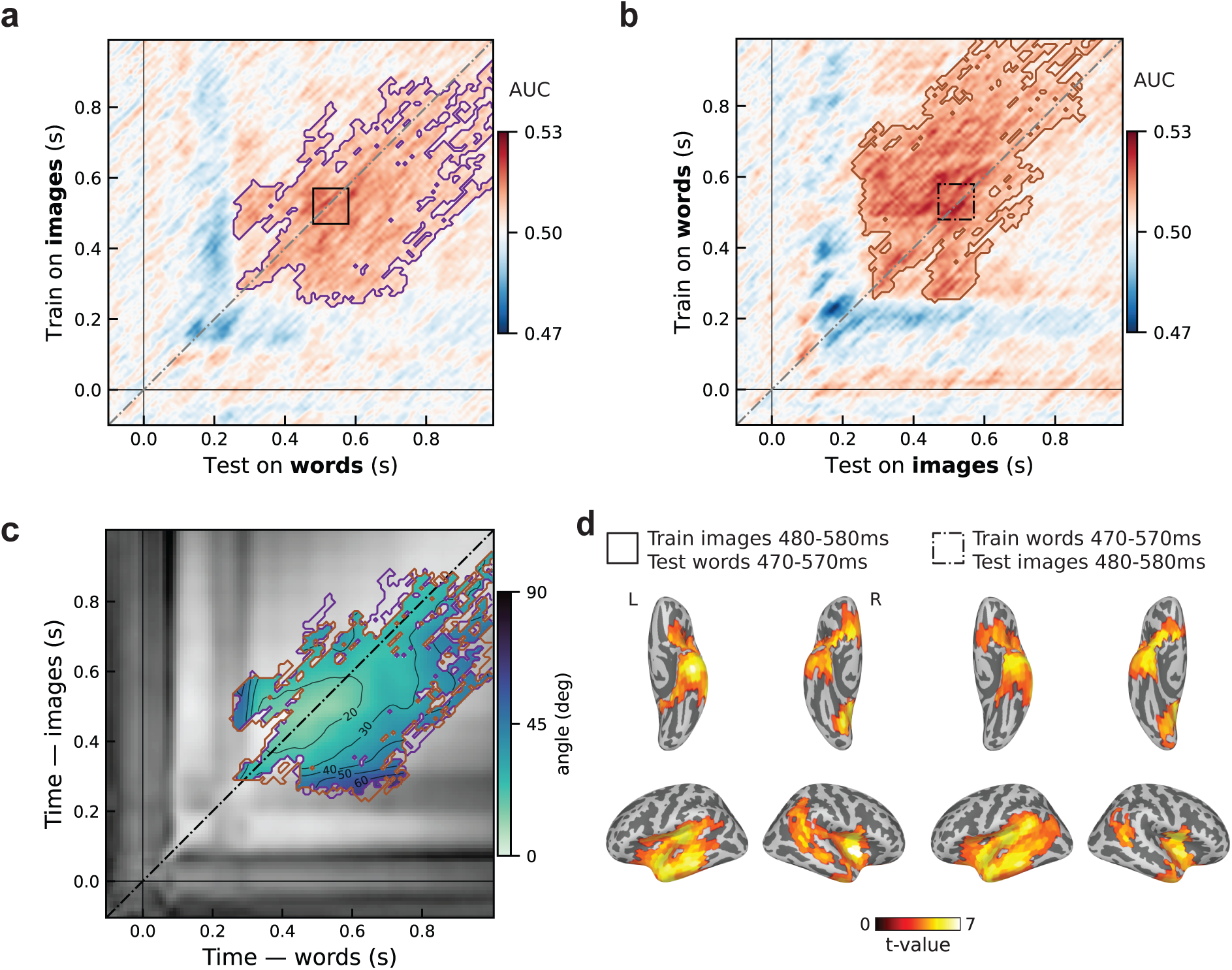
Spatiotemporal characterization of cross-modal representations. (a, b) Temporal generalization matrices for classifiers trained on one modality and tested on the other: trained on images, tested on words (left), and trained on words, tested on images (right). Colour indicates classification performance (AUC) at each training/testing time-point combination. Coloured outlines indicate significant clusters (cluster-based permutation test). Thin black lines mark stimulus onset; the diagonal grey line marks equal training and testing times. (c) Rotation angle between the image and word decision subspaces at every combination of image time (y-axis) and word time (x-axis), averaged across categories. Colour indicates the angle (0° = aligned, 90° = orthogonal), and contour lines mark steps of 20°. The angle is shown only within the significant cross-modal clusters of panels a and b (purple and brown outlines, respectively), with the word-to-image cluster transposed for illustration purposes so that both are plotted with images on the y axis and words on the x axis. The greyscale background shows the angle outside these clusters. The dashed-dotted line marks equal image and word times. (d) Cortical sources of cross-modal generalization, shown as t-values against chance (AUC = 0.5) for significant category-average vertices (cluster-based permutation test). Source-space decoding was performed with a surface searchlight within the 100 ms × 100 ms tiles marked in panels a and b: training on images at 480-580 ms and testing on words at 470-570 ms (solid line in top left panel), and training on words at 470-570 ms and testing on images at 480-580 ms (dash-dotted line in top right panel). Ventral (top row) and lateral (bottom row) views are shown for the left (L) and right (R) hemispheres.

### Relative timing of reaching modality-independent representations in the two modalities

The position of the cross-modal clusters relative to the diagonal indicates which modality reaches the modality-independent representational content first. When training was performed on images and testing on words, the generalization matrix revealed one significant cluster extending primarily below the diagonal: a model trained on images at a given time step generalizes mostly to later time steps in words (Fig. 3a). Training on words and testing on images also revealed a single main cluster, located predominantly above the diagonal: a model trained on words at a given time step generalizes mostly to early time steps of the images (Fig. 3b). This positioning relative to the diagonal describes the net temporal direction of cross-modal generalization: both cross-modal generalization patterns indicate that modality-independent neural representations are reached earlier when items from different semantic categories are presented as images compared to their written-word equivalents.

This observation was confirmed by the diagonal offset index computed over the significant clusters: positive when training on words (0.45), reflecting the predominance of the above-diagonal pattern, and negative when training on images (-0.39), reflecting the predominance of the below-diagonal pattern. The clusters from the two training directions were very similar and appeared mirrored across the diagonal. To compare them directly, we expressed both in a common frame, with image time on the y axis and word time on the x axis, which required transposing the words-to-images matrix. In this frame, 89% of the images-to-words cluster was contained in the words-to-images cluster, and 87% of the words-to-images cluster was contained in the images-to-words cluster (Supplementary Fig. 2). The two clusters thus occupied largely the same region of image time by word time.

Because of this similarity, we located the peak of cross-modal generalization by averaging the two matrices after transposition, which places both in the common frame of image time by word time instead of training time by testing time. The maximum (AUC = 0.52) was at image time 530 ms and word time 520 ms.

### Decision subspace angle

Cross-modal generalization demonstrates shared representations between images and words. However, the generalization performance was weaker than the within-modality performance (peak AUC of 0.52, vs 0.78 for images and 0.59 for words). To test whether this partial generalization rests on aligned decision subspaces (i.e., similar sensor patterns), we computed the angle θ between the image and word decision subspaces for every pair of image and word time points (Fig. 3c, see Supplementary Fig. 3 for within-modality alignment). Angles fell below 20° for image times of approximately 370-590 ms and word times of 330-630 ms, within the window of significant cross-modal generalization. Hence, while cross-modal decoding performance was relatively weak, though reliably above chance, the angle analysis shows that it nonetheless rests on decision subspaces that are largely aligned.

This alignment both strengthens the evidence for a modality-independent code and reinforces the temporal differences across modalities: the aligned window was shorter in image time (220 ms) than in word time (300 ms). Therefore, a shorter period of image time was aligned with a longer period of word time, and the two modalities did not track one another point for point.

### Cross-modal generalization in source space

Sensor-level analyses reported above cannot pinpoint *where* in the cortex the modality-independent code resides, nor whether the two modalities converge on the same regions. We addressed both with a searchlight decoding analysis in source space.

Source-space decoding was applied to the 100 ms × 100 ms tile centred on the peak cross-modal generalization (image time 530 ms, word time 520 ms), a searchlight per cortical vertex. In both directions, classifiers trained on one modality generalized to the other in several significant spatial clusters of the class-averaged AUC maps (images to words: 5 clusters, 480 vertices; words to images: 4 clusters, 414 vertices; Fig. 3d). The clusters of the two directions covered highly similar regions: the bilateral ATL, the bilateral perisylvian cortex (with a larger extent in the left hemisphere), the right inferior parietal cortex, part of the right visual cortex and the bilateral orbitofrontal cortex. The peak AUC was located in the left posterior perisylvian cortex in both directions (images to words: AUC = 0.520, t(31) = 5.50; words to images: AUC = 0.522, t(31) = 5.87; cluster-level p < 0.001).

## Discussion

Our cross-modal decoding findings demonstrate that images and written words converged on a common semantic code within a quarter of a second after stimulus onset (Fig. 3a, b). Unlike previous studies that used tasks such as picture-naming and reading [14], cross-modal matching [13], or semantic judgment [15,16], our task did not refer to either semantic category or modality of the stimulus. Importantly, in the present study each item had to be identified and held in association with the preceding items of its sequence. Thus, the maintaining of a representation was required over a longer interval, whereas in a naming, reading, matching or judgment task the item can be released once a response has been selected. Accordingly, the cross-modal generalization effect was also sustained for longer than in comparable studies, remaining significant for approximately 750 ms within a cluster extending up to 1000 ms, whereas cross-modal generalization lasted for 500 ms [13,15], 400 ms [14] and 300 ms [16] in other studies. The present design placed a demand on the stimulus representation that these that the other designs did not. Another explanation is that the sustained effect is there in other paradigms as well, but our superior decoding performance allows us to detect it.

Cross-modal generalization was asymmetric: classifiers trained on images generalized mostly to later word times, and classifiers trained on words generalized mostly to earlier image times. Asymmetries have been reported in some earlier MEG studies of cross-modal generalization, with one modality reaching the modality-independent representation before the other. Images reached the modality-independent representation first in Leonardelli et al. [16], Bezsudnova et al. [13] and Dirani and Pylkkänen [15]. In Dirani and Pylkkänen [14], generalization first occurred simultaneously for the two modalities (∼125 ms) and at a later stage earlier word states generalized to later image states. Bezsudnova et al. [13] attributed this variability across studies to task demands and designs, such as the presentation of a single category per block in Dirani and Pylkkänen [14], which made the category of each upcoming item predictable and could have led to earlier category access. In the present study, the direction of asymmetry indicated that images reached the modality-independent representation earlier. The finding that an early state of the neural activity elicited by images generalized to a later state of the neural activity elicited by words reinforces the claim of early availability of superordinate category information from the visual form of an image [30–32], while access to the semantic category of a written word necessitates lexical identification of that word [33,34].

The decision subspace analysis gives a more nuanced picture, however. The lowest rotation angles were located on the diagonal, indicating that the decision subspaces were most closely aligned at similar times in the two modalities, around 500 ms (Fig. 3c). The angle analysis also clarifies the origin of the asymmetry in timing between the images-to-words and the words-to-images clusters. The region of low angle was longer on the word axis (300 ms) than on the image axis (220 ms), so the aligned representation was expressed over a longer period when it was elicited by words than by images. Our data indicate that the modalities differ in the direction of the temporal asymmetry of cross-modal generalization and in the duration of the aligned representation, whereas the peak of cross-modal generalization and the region of best alignment occurred at similar times in the two modalities.

Generalization from images to words and from words to images could in principle rely on different signals, for example if each classifier exploited a different component of the other modality’s data. Three findings convergingly indicate that cross-modal generalization reflects a common representation accessed in both modalities. First, the two generalization clusters had nearly the same shape once plotted on the image-time and word-time axes (Supplementary Fig. 2). Thus, the same region of image-time by word-time space was at play whichever modality served as the training set. Second, the decision subspaces of the image and word classifiers were closely aligned for image and word time around 500 ms. In this window, the MEG sensors are weighted similarly by the two classifiers, indicating that category information has a similar expression in the sensor space across modalities. Third, the source-space searchlight yielded significant clusters both when training on images and testing on words and when training on words and testing on images. These clusters were localized largely in the same regions: the bilateral anterior temporal lobe, the perisylvian, visual and orbitofrontal cortex. Together, these results localize the modality-independent code to a selective set of cortical regions, regardless of whether the modality-independent representation is accessed through an image or a written word.

At the peak of cross-modal generalization at ∼500 ms (Fig. 3d), the significant effect extended over regions that correspond broadly to the distributed semantic network described in fMRI studies of semantic processing for words and pictures. In the perisylvian cortex, where the effect was more extensive in the left hemisphere, significant vertices were located in the superior temporal sulcus, the superior and middle temporal gyri and the angular gyrus. These regions have been reported to carry semantic information for both words and images [6–8]. Cross-modal generalization was also significant in the bilateral anterior temporal lobes, consistent with the proposed role of this region as an amodal semantic hub [4,35,36]. Source-space decoding of semantic category from written words alone previously identified the left and right anterior temporal lobe and posterior temporal cortex between approximately 250 and 500 ms [37]. The modality-independent category code in visual cortex is also consistent with Dirani and Pylkkänen [15], who showed that these types of elaborate representations encode visual features. The distribution of the code across the cortex is consistent with a rich representation in which different regions carry different information about the same concept, in line with the semantic selectivity reported across large parts of the cortex with fMRI [38].

### Limitations

The interpretation of these findings is subject to the following limitations. First, category information in the word modality could be carried in part by low-level visual features of the text as a different font was used for each category of word stimuli; furthermore, there was variation in word length across categories. Although this may contribute to the earlier onset of within-modality word decoding than in previous reports, this does not affect the cross-modal findings, as font features are specific to the word modality and cannot support generalization to images. Second, although the task did not require categorization or cross-modal matching, learning and recalling the sequences likely encouraged participants to encode the items at a conceptual level. As each sequence was presented in both modalities, a representation shared by the two modalities could support task performance and thus reinforce modality-independent representations compared with passive viewing of the same stimuli. Third, the class-averaged results may obscure variability in cross-modal generalization across individual semantic categories (Supplementary Fig. 1).

Finally, the source-space analysis has two limitations. In contrast with the temporal decoding analyses based on all sensors, the searchlight was applied separately for each vertex. Hence, it is closer to a mass-univariate approach than to a multivariate one: it captures local patterns but does not capture information distributed across distant regions nor how the regions jointly contribute to the decoding. In addition, source analysis was restricted to the cortical surface, so medial temporal structures that may carry category specific information [39–41] could not be assessed, and given the spatial spread of the inverse solution, part of the signal attributed to the temporal cortex may originate from these deeper sources.

## Conclusion

Images and written words converged on a modality-independent representation of semantic category under a task that referred neither to category membership nor to stimulus modality. Category information was available within each modality from around 70 ms but became transferable between them only from approximately 240 ms, remained so until close to 1000 ms. Activity elicited by images generalized mostly to later activity elicited by words, and a given period of the image response shared a category code with a longer period of the word response. Source estimates place the modality-independent code in a distributed set of cortical regions comprising the perisylvian, orbitofrontal and visual cortex as well as the bilateral anterior temporal lobes, connecting the timing of modality-independent semantic access to the anatomy that functional imaging has previously associated with it (although without its exact timing). Convergence on a modality-independent semantic code appears to be a default of stimulus processing route rather than a product of task demands.

## Acknowledgements

This study was supported by the M/EEG and Neuromod Platform, Fondation Campus Biotech Geneva, Geneva, Switzerland. The authors thank Manon Briet for her help with data collection.

## Author contributions

N.P., T.D., I.O., S.S. and N.K. designed the experiment. N.P. collected the data. N.P. and T.D. wrote the analysis code. N.P. analyzed and interpreted the data. T.D., S.S. and N.K. contributed to data interpretation. N.P. wrote the paper with input from T.D., I.O., S.S. and N.K.

## Competing interest

The authors declare no competing interests.

## Funding sources

This project was funded by the NCCR Evolving Language, Swiss National Science Foundation Agreement, grand number: **#51NF40_225146**.

## Supplementary material

**Supplementary Figure 1.**
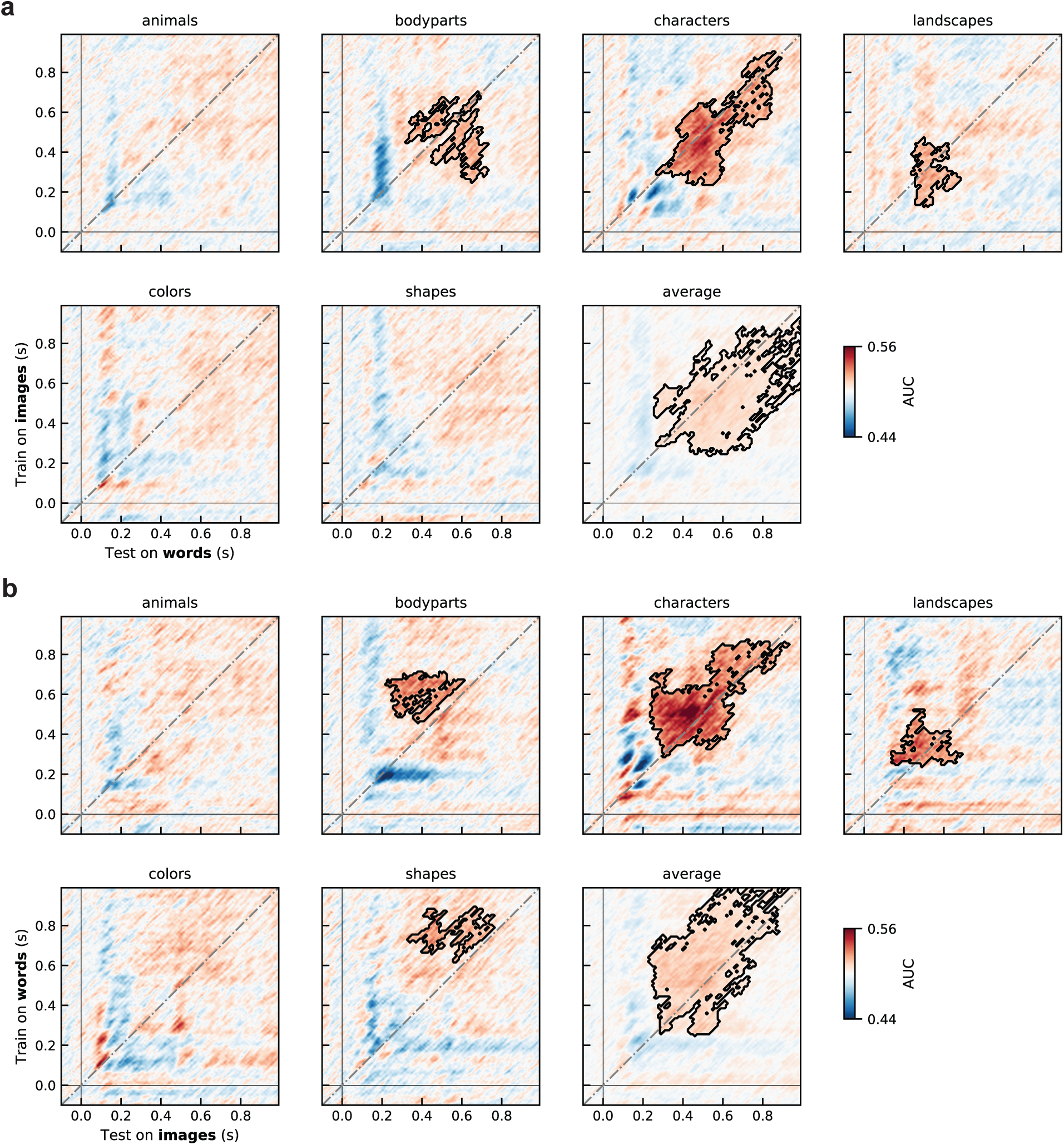
Category breakdown of cross-modal generalization. Cross-modal temporal generalization matrices for individual semantic categories, trained on words and tested on images (a), and trained on images and tested on words (b). Colour indicates classification performance (AUC) at each training/testing time-point combination. Black outlines indicate significant clusters (cluster-based permutation test). Thin black lines mark stimulus onset; the diagonal grey line marks equal training and testing times. When considering categories separately (i.e., reduced number of samples), characters and landscapes still generalized in both directions, body parts and shapes only from words to images, while no significant cross-modality generalization cluster was found for animals and colours.

**Supplementary Figure 2.**
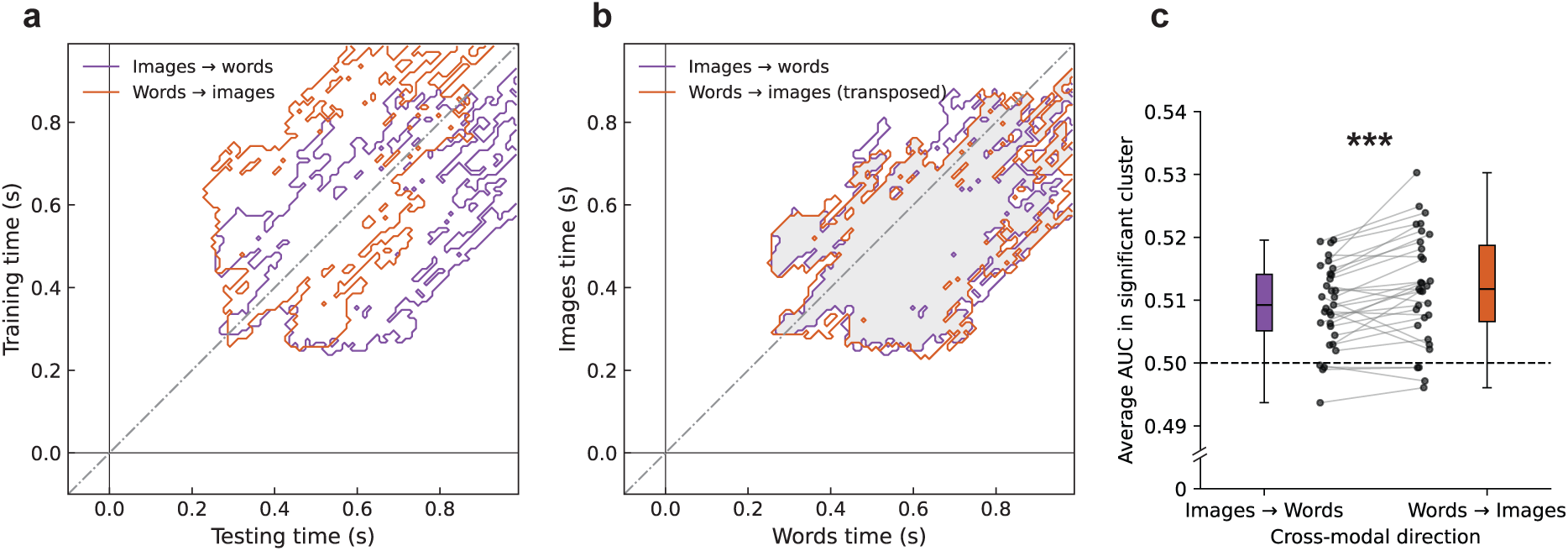
Comparison of cross-modal generalization between the two training directions. (a) Significant cross-modal generalization clusters (cluster-based permutation test) for classifiers trained on images and tested on words (purple) and trained on words and tested on images (orange), plotted on training-time (y-axis) and testing-time (x-axis). The dashed-dotted line marks equal training and testing times. (b) The same clusters replotted on the modality axes, with image time on the y axis and word time on the x axis. The words-to-images cluster is transposed for illustration purposes. The grey area marks the overlap between the two clusters. (c) Average AUC within the significant cluster for each cross-modal direction. Dots show individual participants. The dashed line marks chance (AUC = 0.5). The average generalization is higher when training on words than on images (paired t-test, t(31) = 3.61, p = 0.001).

**Supplementary Figure 3.**
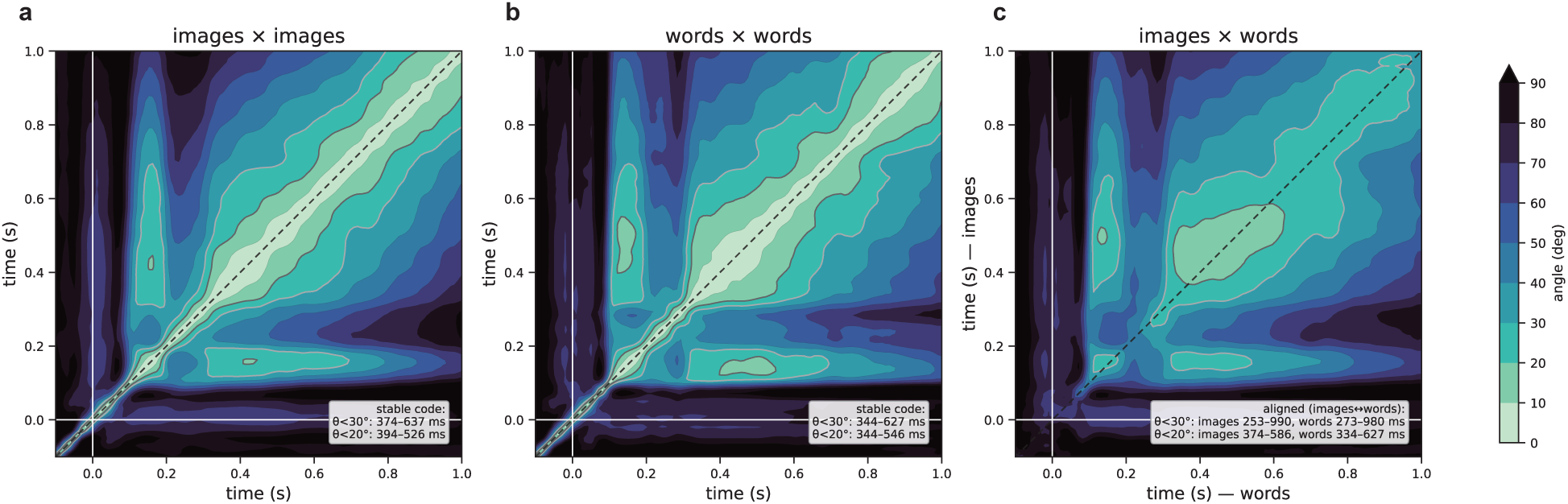
Angles between decision subspaces within and across modalities. Each panel shows the angle θ between activation patterns for every pair of timepoints for images (a), words (b), and cross-modal (c). Matrices were obtained by averaging patterns across the 32 participants before computing θ. Contours were lightly smoothed (Gaussian σ=1 timepoint) for display. Text insets give the well-aligned windows at two shared-variance levels. Because the angle is defined from the normalised dot product of the two pattern vectors (θ = across of their cosine similarity), the cosine of the angle is their correlation and its square, cos²θ, is the proportion of one pattern’s variance captured by the shared direction. We therefore express the alignment thresholds as shared variance: cos²θ ≥ 75% (equivalently θ ≤ 30°) and ≥ 90% (θ ≤ 18°). Brighter colours indicate smaller angles, i.e. more closely aligned subspaces; contours are drawn every 10°. Angles were computed on the 72 sensors retained by the contributing-sensor selection. The diagonal dashed line marks equal latency. White lines mark stimulus onset.

---

**Supplementary Table 1.** Word lengths across categories.

|  | Animals | Body parts | Characters | Colours | Landscapes | Shapes |
| --- | --- | --- | --- | --- | --- | --- |
| Mean | 5.08 | 4.83 | 5.75 | 4.67 | 5.75 | 6.58 |
| SD | 1.19 | 1.21 | 1.01 | 0.74 | 1.48 | 1.44 |

## Notes

### Competing Interest Statement

The authors have declared no competing interest.

## References

1. O’Sullivan FM, Ryan TJ. If Engrams Are the Answer, What Is the Question? In: Gräff J, Ramirez S, editors. Engrams: A Window into the Memory Trace. Cham: Springer International Publishing; 2024. pp. 273–302. doi:10.1007/978-3-031-62983-9_15

2. Poeppel D, Idsardi W. We don’t know how the brain stores anything, let alone words. Trends Cogn Sci. 2022;26: 1054–1055. doi:10.1016/j.tics.2022.08.010

3. Patterson K, Nestor PJ, Rogers TT. Where do you know what you know? The representation of semantic knowledge in the human brain. Nat Rev Neurosci. 2007;8: 976–987. doi:10.1038/nrn2277

4. Lambon Ralph MA, Jefferies E, Patterson K, Rogers TT. The neural and computational bases of semantic cognition. Nat Rev Neurosci. 2017;18: 42–55. doi:10.1038/nrn.2016.150

5. Humphreys GF, Hoffman P, Visser M, Binney RJ, Lambon Ralph MA. Establishing task-and modality-dependent dissociations between the semantic and default mode networks. Proc Natl Acad Sci. 2015;112: 7857–7862. doi:10.1073/pnas.1422760112

6. Humphreys GF, Lambon Ralph MA. A lateral temporal network for transmodal combinatorial semantics: Convergent evidence from a multi-study investigation. bioRxiv; 2026. p. 2026.02.19.706785. doi:10.64898/2026.02.19.706785

7. Devereux BJ, Clarke A, Marouchos A, Tyler LK. Representational Similarity Analysis Reveals Commonalities and Differences in the Semantic Processing of Words and Objects. J Neurosci. 2013;33: 18906–18916. doi:10.1523/JNEUROSCI.3809-13.2013

8. Fairhall SL, Caramazza A. Brain Regions That Represent Amodal Conceptual Knowledge. J Neurosci. 2013;33: 10552–10558. doi:10.1523/JNEUROSCI.0051-13.2013

9. Kaplan JT, Man K, Greening SG. Multivariate cross-classification: applying machine learning techniques to characterize abstraction in neural representations. Front Hum Neurosci. 2015;9. doi:10.3389/fnhum.2015.00151

10. Meyer K, Kaplan JT, Essex R, Webber C, Damasio H, Damasio A. Predicting visual stimuli on the basis of activity in auditory cortices. Nat Neurosci. 2010;13: 667–668. doi:10.1038/nn.2533

11. Simanova I, Gerven M van, Oostenveld R, Hagoort P. Identifying Object Categories from Event-Related EEG: Toward Decoding of Conceptual Representations. PLOS ONE. 2010;5: e14465. doi:10.1371/journal.pone.0014465

12. King J-R, Dehaene S. Characterizing the dynamics of mental representations: the temporal generalization method. Trends Cogn Sci. 2014;18: 203–210. doi:10.1016/j.tics.2014.01.002

13. Bezsudnova Y, Quinn AJ, Wynn SC, Jensen O. Spatiotemporal Properties of Common Semantic Categories for Words and Pictures. J Cogn Neurosci. 2024;36: 1760–1769. doi:10.1162/jocn_a_02182

14. Dirani J, Pylkkänen L. The time course of cross-modal representations of conceptual categories. NeuroImage. 2023;277: 120254. doi:10.1016/j.neuroimage.2023.120254

15. Dirani J, Pylkkänen L. MEG Evidence That Modality-Independent Conceptual Representations Contain Semantic and Visual Features. J Neurosci. 2024;44. doi:10.1523/JNEUROSCI.0326-24.2024

16. Leonardelli E, Fait E, Fairhall SL. Temporal dynamics of access to amodal representations of category-level conceptual information. Sci Rep. 2019;9: 239. doi:10.1038/s41598-018-37429-2

17. Peirce J, Gray JR, Simpson S, MacAskill M, Höchenberger R, Sogo H, et al. PsychoPy2: Experiments in behavior made easy. Behav Res Methods. 2019;51: 195–203. doi:10.3758/s13428-018-01193-y

18. Gramfort A, Luessi M, Larson E, Engemann DA, Strohmeier D, Brodbeck C, et al. MEG and EEG data analysis with MNE-Python. Front Neurosci. 2013;7. doi:10.3389/fnins.2013.00267

19. Taulu S, Simola J. Spatiotemporal signal space separation method for rejecting nearby interference in MEG measurements. Phys Med Biol. 2006;51: 1759. doi:10.1088/0031-9155/51/7/008

20. Pedregosa F, Varoquaux G, Gramfort A, Michel V, Thirion B, Grisel O, et al. Scikit-learn: Machine Learning in Python. J Mach Learn Res. 2011 [cited 29 July 2026]. Available: https://inria.hal.science/hal-00650905

21. Desbordes T, Olasagasti I, Piron N, Schwartz S, Kazanina N. Temporal evolution of Neural Codes: The Added Value of a Geometric Approach to Linear Coefficients. NeuroImage. 2026; 121737. doi:10.1016/j.neuroimage.2026.121737

22. Haufe S, Meinecke F, Görgen K, Dähne S, Haynes J-D, Blankertz B, et al. On the interpretation of weight vectors of linear models in multivariate neuroimaging. NeuroImage. 2014;87: 96–110. doi:10.1016/j.neuroimage.2013.10.067

23. Krogh SM, Pylkkänen L. Comparing serial and parallel language input: Shared neural representations with distinct encoding strengths. bioRxiv; 2026. p. 2026.09.23.753812. doi:10.64898/2026.09.23.753812

24. Zuanazzi A, Ripollés P, Lin WM, Gwilliams L, King J-R, Poeppel D. Negation mitigates rather than inverts the neural representations of adjectives. PLOS Biol. 2024;22: e3002622. doi:10.1371/journal.pbio.3002622

25. Maris E, Oostenveld R. Nonparametric statistical testing of EEG- and MEG-data. J Neurosci Methods. 2007;164: 177–190. doi:10.1016/j.jneumeth.2007.03.024

26. Sassenhagen J, Draschkow D. Cluster-based permutation tests of MEG/EEG data do not establish significance of effect latency or location. Psychophysiology. 2019;56: e13335. doi:10.1111/psyp.13335

27. Destrieux C, Fischl B, Dale A, Halgren E. Automatic parcellation of human cortical gyri and sulci using standard anatomical nomenclature. NeuroImage. 2010;53: 1–15. doi:10.1016/j.neuroimage.2010.06.010

28. Ghazaryan G, van Vliet M, Saranpää A, Lammi L, Lindh-Knuutila T, Hultén A, et al. Trials and tribulations when attempting to decode semantic representations from MEG responses to written text. Lang Cogn Neurosci. 2024;39: 1149–1160. doi:10.1080/23273798.2023.2219353

29. Rybář M, Daly I. Neural decoding of semantic concepts: a systematic literature review. J Neural Eng. 2022;19: 021002. doi:10.1088/1741-2552/ac619a

30. Fabre-Thorpe M. The Characteristics and Limits of Rapid Visual Categorization. Front Psychol. 2011;2: 243. doi:10.3389/fpsyg.2011.00243

31. Macé MJ-M, Joubert OR, Nespoulous J-L, Fabre-Thorpe M. The Time-Course of Visual Categorizations: You Spot the Animal Faster than the Bird. PLOS ONE. 2009;4: e5927. doi:10.1371/journal.pone.0005927

32. Wu C-T, Crouzet SM, Thorpe SJ, Fabre-Thorpe M. At 120 msec You Can Spot the Animal but You Don’t Yet Know It’s a Dog. J Cogn Neurosci. 2015;27: 141–149. doi:10.1162/jocn_a_00701

33. Coltheart M, Rastle K, Perry C, Langdon R, Ziegler J. DRC: A dual route cascaded model of visual word recognition and reading aloud. Psychol Rev. 2001;108: 204–256. doi:10.1037/0033-295X.108.1.204

34. Hauk O, Davis MH, Ford M, Pulvermüller F, Marslen-Wilson WD. The time course of visual word recognition as revealed by linear regression analysis of ERP data. NeuroImage. 2006;30: 1383–1400. doi:10.1016/j.neuroimage.2005.11.048

35. Mesulam MM. From sensation to cognition. Brain J Neurol. 1998;121 ( Pt 6): 1013–1052. doi:10.1093/brain/121.6.1013

36. Mesulam M-M, Wieneke C, Hurley R, Rademaker A, Thompson CK, Weintraub S, et al. Words and objects at the tip of the left temporal lobe in primary progressive aphasia. Brain. 2013;136: 601–618. doi:10.1093/brain/aws336

37. Magnabosco F, Hauk O. Decoding Semantics from Dynamic Brain Activation Patterns: From Trials to Task in EEG/MEG Source Space. eNeuro. 2024;11. doi:10.1523/ENEURO.0277-23.2023

38. Huth AG, de Heer WA, Griffiths TL, Theunissen FE, Gallant JL. Natural speech reveals the semantic maps that tile human cerebral cortex. Nature. 2016;532: 453–458. doi:10.1038/nature17637

39. Heusser AC, Poeppel D, Ezzyat Y, Davachi L. Episodic sequence memory is supported by a theta–gamma phase code. Nat Neurosci. 2016;19: 1374–1380. doi:10.1038/nn.4374

40. Tacikowski P, Kalender G, Ciliberti D, Fried I. Human hippocampal and entorhinal neurons encode the temporal structure of experience. Nature. 2024;635: 160–167. doi:10.1038/s41586-024-07973-1

41. Wimmer GE, Liu Y, McNamee DC, Dolan RJ. Distinct replay signatures for prospective decision-making and memory preservation. Proc Natl Acad Sci U S A. 2023;120: e2205211120. doi:10.1073/pnas.2205211120

